# Targeting the chemokine receptor CXCR4 with histamine analogue to reduce inflammation in juvenile arthritis: a proof of concept for COVID-19 therapeutic approach

**DOI:** 10.1101/2021.10.24.465080

**Authors:** Nassima Bekaddour, Nikaïa Smith, Benoit Beitz, Alba Llibre, Tom Dott, Anne Baudry, Anne-Sophie Korganow, Sébastien Nisole, Richard Mouy, Sylvain Breton, Brigitte Bader-Meunier, Darragh Duffy, Benjamin Terrier, Benoit Schneider, Pierre Quartier, Mathieu P Rodero, Jean-Philippe Herbeuval

## Abstract

Among immune cells, activated monocytes play a detrimental role in chronic and viral-induced inflammatory pathologies. The uncontrolled activation of monocytes and the subsequent excessive production of inflammatory factors damage bone-cartilage joints in Juvenile Idiopathic Arthritis (JIA), a childhood rheumatoid arthritis (RA) disease. Inflammatory monocytes also exert a critical role in the cytokine storm induced by SARS-CoV2 infection in severe COVID-19 patients. The moderate beneficial effect of current therapies and clinical trials highlights the need of alternative strategies targeting monocytes to treat RA and COVID-19 pathologies. Here, we show that targeting CXCR4 with small amino compound such as the histamine analogue clobenpropit (CB) inhibits spontaneous and induced-production of a set of key inflammatory cytokines by monocytes isolated from blood and synovial fluids of JIA patients. Moreover, daily intraperitoneal CB treatment of arthritic mice results in significant decrease in circulating inflammatory cytokine levels, immune cell infiltrates, joints erosion, and bone resorption leading to reduction of disease progression. Finally, we provide the prime evidence that the exposure of whole blood from hospitalized COVID-19 patients to CB significantly reduces levels of key cytokine-storm-associated factors including TNF-α, IL-6 and IL-1β. These overall data show that targeting CXCR4 with CB-like molecules may represent a promising therapeutic option for chronic and viral-induced inflammatory diseases.

## INTRODUCTION

Rheumatoid arthritis (RA) is a chronic inflammatory disease associated with aggressive synovial hyperplasia causing destruction of articular joints. Joint inflammation is characterized by proliferation of macrophage-like [1] and fibroblast-like synoviocytes to form a pannus, which invades and destroys the cartilage [2]. Juvenile Idiopathic Arthritis (JIA) is a rare and complex form of RA occurring in children under the age of 16, characterized by a multifactorial disorder with heterogeneous manifestations that include all forms of chronic arthritis [3,4]. Over-production of TNF-α, IL-1β and IL-6 is strongly involved in most forms of JIA [5][6]. The release of such inflammatory factors by monocytes depends on several highly conserved families of pattern recognition receptors (PRR), one of them being the Toll-like receptor family (TLRs). The increase in circulating TNF-α levels in all forms of JIA argues for a major contribution of monocytes to disease progression [7]. The role of monocytes in JIA has been further highlighted in a work showing that stimulation of TLR-4 and TLR-8 in monocytes isolated from the blood of systemic JIA patients increased the proinflammatory responses compared to healthy donors [8]. Current antirheumatic drugs, including corticosteroids and antibodies-based biotherapies, target inflammatory macrophages to reduce synovial inflammation. However, not all patients respond to antibody therapy and chronic application of glucocorticoids leads to severe side effects, highlighting the need for novel therapeutic strategies.

Monocytes are also suspected to play a deleterious role during acute inflammation upon SARS-CoV-2 infection responsible for Coronavirus disease 2019 (COVID-19). Indeed, SARS-CoV2 infects the upper and lower respiratory tracts and causes mild or highly acute respiratory syndrome with consequent release of proinflammatory cytokines by hyper-activated monocytes [9,10]. Recently, we measured over-production of proinflammatory cytokines in COVID-19 patients causing cytokine storm in lungs[10]. Furthermore, high levels of IL-6 and TNF-α in bronchoalveolar lavage and plasma of those patients corelate with the severity of the pathology [10]. This SARS-CoV2 pandemic stressed the scientific and clinical communities to urgently develop novel anti-inflammatory strategies.

In the context of chronic and acute inflammatory diseases, the chemokine receptor CXCR4 could emerge a potential therapeutic target. We previously showed that histamine and the histamine analogue clobenpropit (CB) through their engagement of CXCR4 exhibited a broad-spectrum inhibitory activity on the production of all subtypes of interferons (IFN) in TLR7-activated human plasmacytoid dendritic cells (pDCs) [11]. Moreover, intranasal spray of CB resulted in drastic reduction of type I and III IFN secretions in broncho-alveolar lavages from Influenza A virus (IAV) infected mice [11]. Of note, the anti-IFN activity of CB was not mediated by histamine receptors but strictly dependent on CXCR4 [11]. As CXCR4 is highly expressed by all immune cells, including monocytes and macrophages, we hypothesize that CB could also downmodulate monocyte-driven inflammation in rheumatoid arthritis and SARS-CoV2 infection.

Here, we assessed whether CB tones down the spontaneous and induced production of proinflammatory cytokines by monocytes isolated from blood and synovial fluids from JIA patients. We further probed *in vivo* whether CB treatment would attenuate inflammation and impair disease progression in a model of collagen-induced arthritic mice. Finally, we tested the effect of CB on the production of inflammatory cytokines in whole blood from hospitalized COVID-19 patients.

## RESULTS

### CB down-regulates TNF-α, IL-1β, and IL-6 productions in TLR-7/8-activated monocytes

To first assess the potential anti-inflammatory properties of histamine and the histamine analogue CB, we took advantage of the monocyte-derived THP-1 NF-κB reporter cell line. Upon TLR7/8 activation by R848, the reporter gene SEAP is induced by activation of the NF-κB signaling pathway. We thus measured the production of SEAP by R848-activated THP-1 cells in the presence or absence of increasing concentration of histamine (**Fig. 1A**) or CB (**Fig. 1B**) ranging from 1 to 100 μM. We showed that CB, and to a lesser extent histamine, reduced in a dose-dependent manner activation of NF-κB in THP-1 cells, without any obvious toxicity. In addition, CB treatment (20 μM) prevented the transcription of TNF-α, IL-1β and IL-6 encoding genes in R848-activated THP-1 cells (**Fig. 1C**).

**Figure 1.**
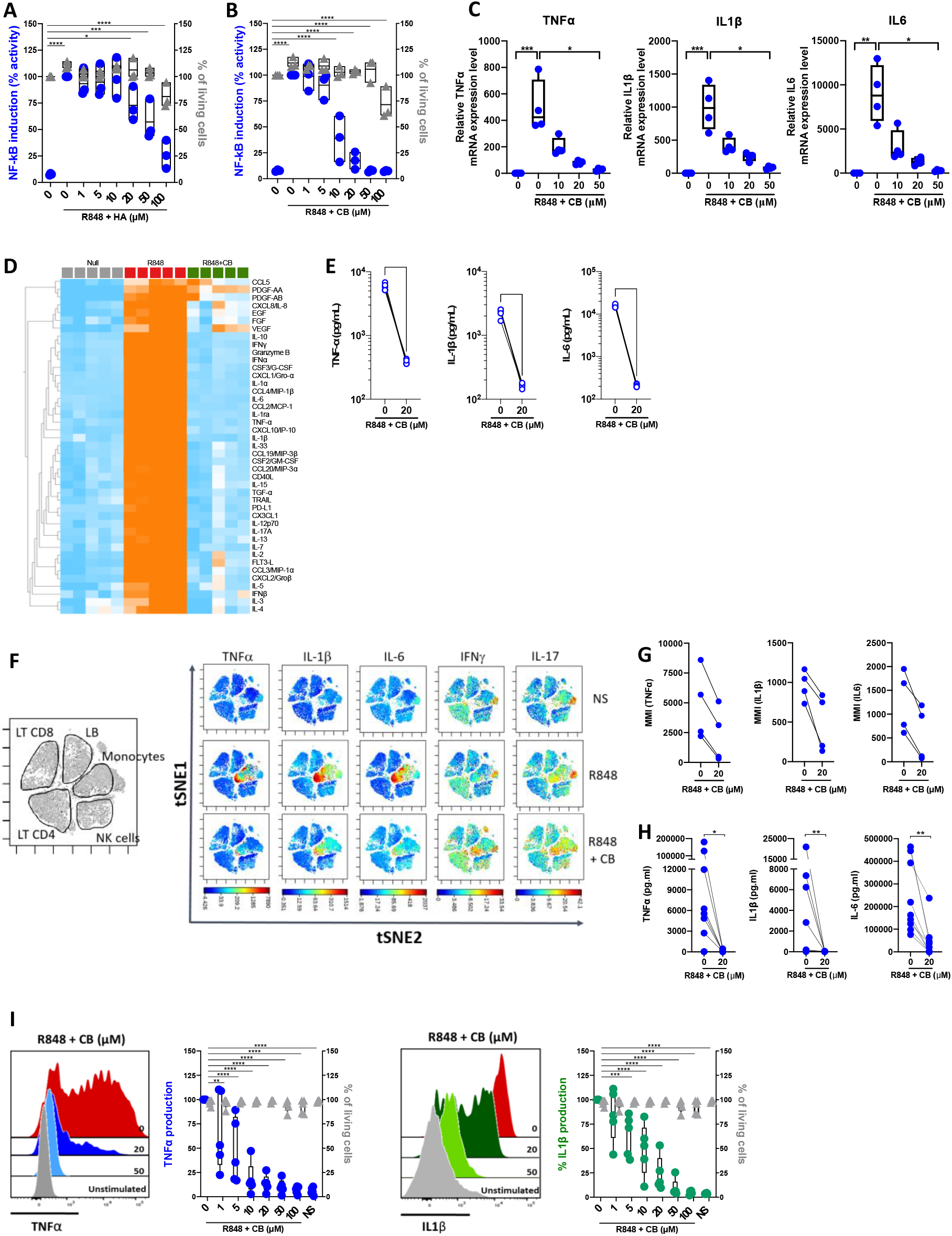
CB inhibited TLR-7/8-mediated TNF-α, IL-1β and IL-6 production by human monocytes. (**A-B**) THP1 NF-κB reporter cells were treated with increasing concentration of Histamine (**A**) or CB (**B**) ranging from 1 to 100 μM then stimulated with R848 (5 μg/ml) for 24 hours. NF-κB reporter activity was measured using QUANTI-Blue, a SEAP detection reagent. (**C**) mRNA levels of TNF-α, IL-1β and IL-6 from THP1-dual pre-incubated with increased doses of CB and stimulated overnight with R848 (5 μg/ml), were measured by RT-qPCR and normalized to RPL13A. Kruskal-Wallis with Dunn’s multiple comparisons test. (**D-G**) PBMCs from healthy donors (HD) were preincubated with CB (20 μM) then stimulated with R848 (5 μg/mL) overnight. (**D-E**) Cytokine production was measured in the supernatant using a bead-based multiplexed immunoassay system Luminex. (**D**) Heatmap representation of statistically different cytokines (P<0.05) between the different conditions (Null, R848, R848+CB), ordered by hierarchical clustering. Up-regulated cytokines are shown in orange, and down-regulated in blue. P values were determined with the Kruskal-Wallis test. (**E**) Individual cytokines from the Luminex are represented. Mann-Whitney test. (**F**) PBMCs were analyzed by mass cytometry (CyTOF) and tSNE analysis were performed using CD56, CD3, CD11c, BDCA4, CD14, HLADR, CD123, CD4, CD8, and CD19 markers. Intracellular levels of TNF-α, IL-1β, IL-6, IFNγ, and IL-17 were evaluated. (**G**) Mean Metal Intensity (MMI) of TNF-α, IL-1β, and IL-6 was evaluated on gated monocytes from four different donors. (**H**) Cytokine production was measured in the supernatants of purified monocytes from nine HD using the multiplex bead-based immunoassay LEGENDplex. Mann-Whitney test. (**I**) Purified monocytes from five HD were preincubated with increased doses of CB then stimulated with R848 during 5 h. Intracellular levels of TNF-α and IL-1β as well as viability were evaluated by flow cytometry. SSC-A, side scatter. FSC-A, forward scatter. 2-way ANOVA. All data are presented as median ± range. ****P < 0.0001, ***P < 0.001, **P < 0.01, *P < 0.05.

To extend our results obtained with THP-1 cell line, we next tested the ability of CB to control cytokine production in R848-stimulated human blood mononuclear cells (PBMCs) from healthy donors (HD). We first tested the overall effect of CB by measuring the concentrations of multiple cytokines (*i.e.* IL-1β, −6, −8, −10, TNF-α…), chemokines (*i.e.* CCL2, CCL3, CCL4, CCL5…), growth factors (*i.e.* EGF, FGF, VEGF…) and interferons (IFNα/β/γ) in the cell culture medium of R848-stimulated PBMCs in presence or absence of CB (20 μM) (n=5) (**Fig. 1D**). CB downmodulated the R848-induced production of chemokines, growth factors, IFN subtypes, and all proinflammatory cytokines, including TNF-α, IL-1β and IL-6 (**Fig. 1D and 1E**). Mass Cytometry (CyTOF) analysis revealed that among PBMCs monocytes represented the main producers of TNF-α, IL-1β and IL-6 upon TLR7/8 stimulation by R848. CB treatment drastically reduced the production of those inflammatory factors by R848-activated monocytes **(Fig. 1F, G)** (n=4). To better evaluated the effect of CB on the proinflammatory cytokines, we purified monocytes from the blood of HD. Accordingly, using a multiplex bead-based assay (**Fig. 1H**) and intracellular flow cytometry staining of TNF-α and IL-1β, we showed in monocytes purified from HD that CB inhibited the production of TNF-α and IL-1β (IC_50_=2,8 μM and IC_50_=8 μM, respectively) with no observed toxicity (**Fig. 1I**).

### CB controls activation monocytes from healthy individuals in a CXCR4-dependent manner

We next tested whether the anti-inflammatory properties of CB depend on CXCR4. We first used the well-known CXCR4 antagonist AMD3100 to modulate the anti-inflammatory effects of CB on activated PBMCs. Cells were cultured with increased concentrations of AMD3100 in presence of CB (20μM) and R848 for 24h. Levels of IFN and pro-inflammatory factors produced by activated PBMC were indirectly quantified using the monocyte-derived IRF and NF-κB reporter cell line THP-1 dual. We showed that CB reduced activation of both IRF and NF-κB and that AMD3100 blocked the anti-inflammatory activity of CB in a concentration dependent manner (**Fig. 2A–B**).

**Figure 2.**
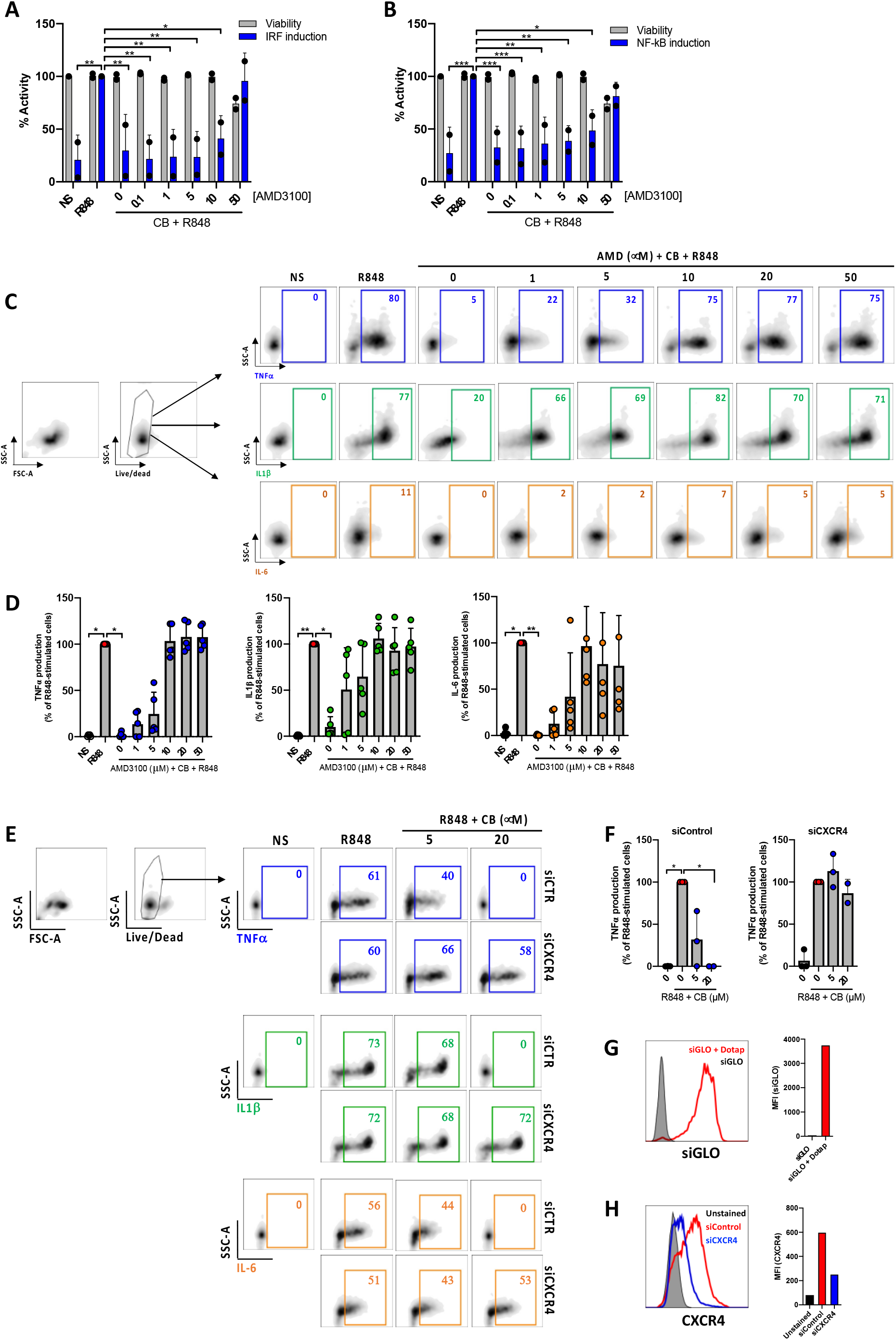
CB immunoregulatory activity on primary monocytes is CXCR4 dependent. (**A**) and (**B**) PBMCs from two healthy donors were cultured in presence of increased concentrations of AMD3100, then treated with CB (20 μM) and activated with R848 (5 μg/mL) for 24 hours. The level of (**A**) interferons and (**B**) inflammatory cytokines in the culture supernatants was measured using the reporter cell line THP1-dual. Two-way ANOVA with Dunnett’s multiple comparisons test. (**C**) and (**D**) Isolated monocytes from five healthy donors were cultured in presence of increased concentrations of AMD3100, then treated with CB (20 μM) and activated with R848 (1 μg/mL) for 6 hours. Intracellular levels of TNF-α, IL-1b and IL-6 were evaluated by flow cytometry, (**C**) dot plot representation from a donor and (**D**) histogram representation from 5 donors. Friedman with Dunn’s multiple comparisons test. (**E-F**) Monocytes were treated for 24 hours with CXCR4 siRNA (siCXCR4) or Control siRNA (siControl) at 160 nM. Cells were then pre-incubated with CB (20 μM) and stimulated with R848 (1μg/mL) for 6 hours. Intracellular levels of TNF-α, IL-1b and IL-6 were evaluated by flow cytometry: (**E**) dot plot representation from a donor and (**F**) histogram representation from 3 donors. Data shown as the median +/-s.d. Krustal-Wallis with Dunn’s multiple comparisons test. (**G**) the efficiency of transfection with dotap on monocytes was evaluated with GLO siRNA in flow cytometry. (**H**) The expression of CXCR4 on monocytes treated with siControl and siCXCR4 was assessed by flow cytometry. **P < 0.01, *P < 0.05.

We further quantified TNF-α, IL1β and IL-6 productions in R848-stimulated primary purified monocytes from HD in the presence of CB (20 μM) and increasing concentrations of AMD3100 (up to 50 μM) using flow cytometry. As expected, AMD3100 inhibited in a dose-dependent manner the ability of CB to reduce intracellular TNF-α, IL1β and IL-6 levels in monocytes without noticeable toxicity (**Fig. 2C–D**).

To firmly establish that CB immunoregulatory activity relates to CXCR4, the expression of CXCR4 was silenced in primary monocytes using small interfering RNA (siRNA). While the transfection of a siRNA control (siControl) had no impact on the CB-induced inhibition of pro-inflammatory cytokines, CB lost its ability to inhibit TNF-α, IL1β and IL-6 intracellular productions in CXCR4-silenced human monocytes (siCXCR4) (**Fig. 2E–F**). Taking into account the 95% transfection efficiency in monocytes, we checked that CXCR4 expression was strongly reduced in siCXCR4-transfected monocytes compared to siControl (**Fig. 2G–H**). These overall data show that the immunoregulatory effects of CB strictly depends on CXCR4 engagement.

### CB inhibits spontaneous proinflammatory cytokine production in JIA patients’ monocytes

Our next attempt was to prob whether CB exerts an anti-inflammatory effect on immune cells isolated from AJI patients. Spontaneous inflammation is a well-known hallmark of JIA. While elevated levels of IL-6 are easily measurable in the plasma of RA patients [12], the detection of circulating TNF-α in these patients remains a challenge. Thus, to capture TNF-α in the plasma, we developed a TNF-α digital ELISA (Simoa) that detects TNF-α for concentrations as low as 1 fg/mL. Using this technology, we measured detectable levels of TNF-α in patients’ plasma and found higher of TNF-α in blood plasma from JIA patients than in HD (JIA patients’ median = 3.4 pg/mL *vs.* HD’ median = 1.07 pg/mL) **(Fig. 3A**). To better capture the spontaneous inflammation in JAI patients, we performed a gene expression array of 579 inflammatory markers on purified monocytes of PBMC from HD and JAI patients. As compared to HD, the expression of 51 inflammatory genes significantly increased and 16 decreased in JIA patients (**Fig. 3B**). Principal components analysis (PCA) showed that monocytes from each group clustered together (**Fig. 3C**). Pathway analysis using DAVID [13,14] showed enrichment in the rheumatoid arthritis pathway (**Fig. 3D**). Since JIA is primarily a joint-related inflammatory disease [15], this prompted us to examine the inflammatory status of synovial fluids. According to Simoa experiments, we measured TNF-α concentrations between 3 and 26 pg/mL (median = 10.8 pg/mL) in synovial fluid (SF) from eighteen JIA patients’ knees with active arthritis (**Fig. 3E**). When comparing circulating levels of TNF-α in plasma and SF from six matched patients, higher levels of TNF-α were systematically measured in SF compared to blood plasma (**Fig. 3F**).

**Figure 3.**
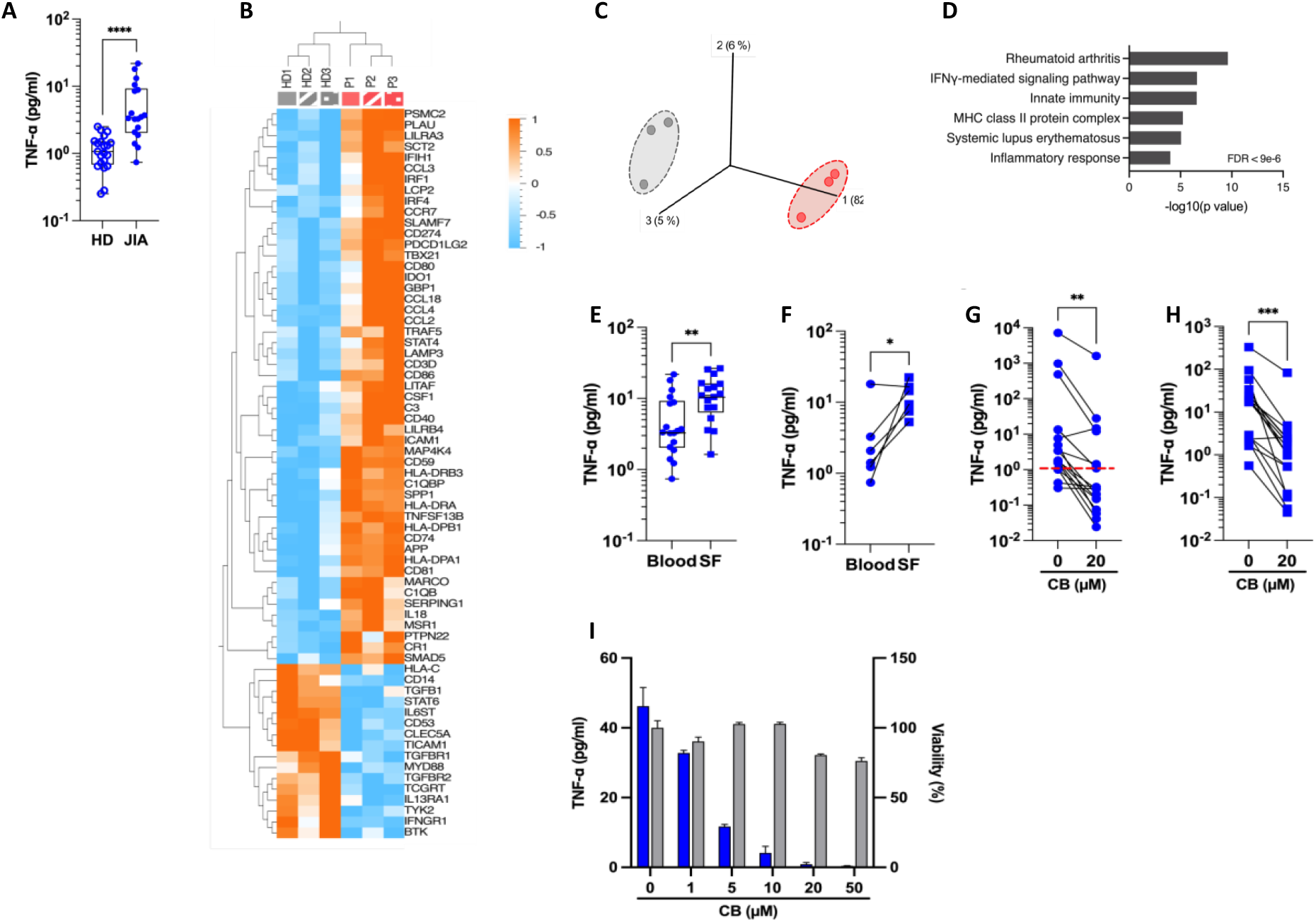
CB controls spontaneous inflammation in cells from JIA patients. (**A**) TNF-α was measured in the plasma of blood from HD and JIA patients by digital ELISA (Simoa). Box and whisker plots with median ± range. Mann-Whitney test. (**B**) Monocytes from blood from healthy donors (HD) and JIA patients were isolated and total RNA from the cells was isolated and analyzed using the NanoString nCounter. Heat map represents mRNA fold increase of the genes significantly differently expressed between the two groups (P < 0.05, fold change > 1.5). (**C**) PCA was performed based on genes from the array differentially expressed between the HD (gray) and the JIA patients (red). (P < 0.05, fold change > 1.5). (**D**) David pathway analysis was used for pathway enrichment analysis of differentially upregulated genes in JIA patients compared to the HD. (**E**) TNF-α was dosed in the plasma from blood or synovial fluids (SF) of JIA patients by Simoa. Mann-Whitney test. (**F**) TNF-α levels in paired plasma and SF from six different donors were dosed by Simoa. Wilcoxon test. TNF-α was measured in the supernatant of monocytes by digital ELISA (Simoa) purified from PBMCs (**G**) or SFMCs (**H**) of JIA patients. Average baseline of HD is represented by the red dashed line. Mann-Whitney test. (**I**) Monocytes from one JIA patient were treated with various concentration of CB for 16h and TNF-α was measured in the supernatant by digital ELISA (Simoa).

We next evaluated the impact of CB treatment on the spontaneous TNF-α production by human monocytes isolated from PBMCs (**Fig. 3G**) and SFMCs (**Fig. 3H**) from JIA patients. CB treatment significantly decreased TNF-α levels in the culture medium of both SFMCs and PBMCs monocytes. We also showed that CB exerted a dose-dependent anti-inflammatory effect in purified monocytes from SF of one patient without any significant toxicity (**Fig. 3I**). Altogether, these data provide evidence that CB inhibits the spontaneous production of inflammatory cytokines by blood and synovial human monocytes within an arthritic context.

### CB attenuates R848-induced cytokine production in JIA patients’ cells

We next evaluated whether CB would also inhibit cytokine production by stimulated immune cells from JAI patients. To this purpose, we first compared cytokine production by PBMCs from HD and JIA patients using the 45-plex bead-based immunoassay Luminex. We showed that PBMCs from three JIA patients out of five secreted more cytokines than those of HD **(Fig. 4A**). This result confirms that cells from JIA patients are hyperresponsive to TLR-7/8 stimulation [8]. Furthermore, CB inhibited R848-induced production of inflammatory cytokines by PBMC from HD as well as the hypersecretion of inflammatory cytokines from JIA patients’ cells (**Fig. 4B**). As monocytes are the main producers of proinflammatory cytokines, we purified monocytes from blood patients’ suffering from oligo articular (n=20), polyarticular (n=8) or systemic (n=5) JIA, or enthesitis-related arthritis (ERA) (n=3). For all subtypes of JIA, CB treatment drastically reduced levels of IL-1β, IL-6, and TNF-α in the culture medium of R848-activated patients’ monocytes (**Fig. 4C**).

**Figure 4.**
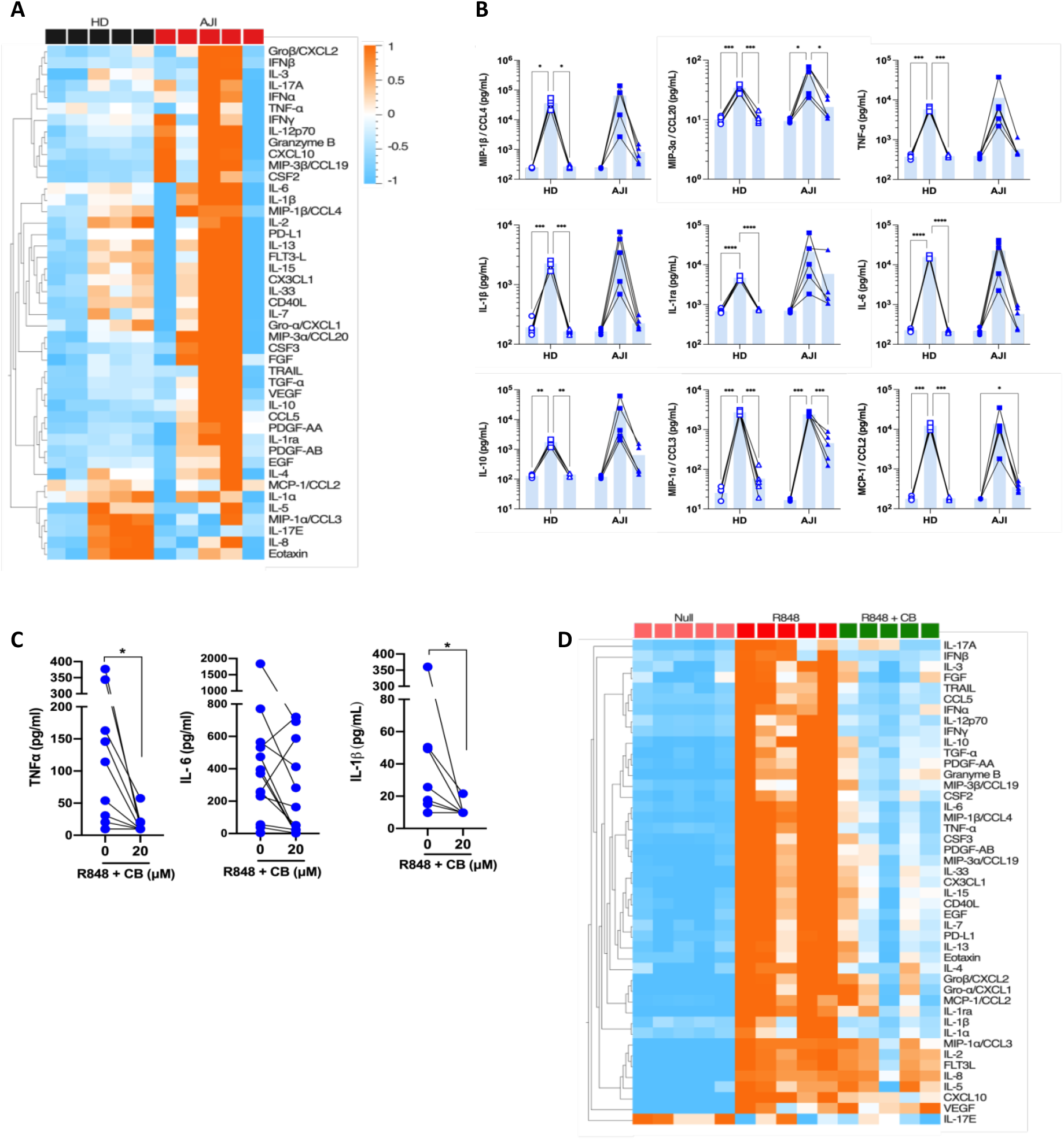
CB controls R848-induced inflammation in cells from JIA patients. (**A**) and (**B**) PBMCs from HD and JIA patients were pre-incubated with CB at 20 μM and then stimulated with R848 (5 μg/mL) for 16h. Cytokine production was measured in the supernatant using a bead-based multiplexed immunoassay system Luminex. (**A**) Heatmap representation of statistically different cytokines (P<0.05) in PBMCs supernatant upon R848 stimulation between HD and JIA patients, ordered by hierarchical clustering. Mann-Whitney test. (**B**) Individual cytokines are represented. Two-way RM ANOVA with Tukey post hoc correction. (**C**) Monocytes from JIA patients were pre-incubated with CB at 20 μM and then stimulated with R848 (5 μg/mL) for 16h. IL-6, IL-1β and TNF-α production was measured in the supernatant using the multiplex bead-based immunoassay LEGENDplex. Mann-Whitney test. (**D**) SFMCs were pre-incubated with CB at 20 μM and then stimulated with R848 (5 μg/mL) for 16h. Cytokine production was measured in the supernatant using a bead-based multiplexed immunoassay system Luminex.

Finally, using purified synovial fluid mononuclear cells (SFMCs) from JIA patients we showed that CB inhibited the production of the 40 soluble factors induced by R848 activation (**Fig. 4D**). Overall data indicate that CB could reduce hyper secretion of inflammatory cytokines observed in JAI patients during flares.

### CB inhibits inflammatory cytokine and reduces disease progression in collagen-induced arthritis mouse model

The anti-inflammatory properties of CB in PBMCs, SFMCs and monocytes of JIA patients, suggest that CB represents a promising therapeutic option for arthritis. Thus, we tested the potential therapeutic effect of CB in a mouse model with an arthritis-like pathology, the collagen-induced arthritis (CIA) DBA/1J mice [16]. CIA mice share many clinical, histologic, and immunologic features with human RA [17]. This includes symmetric joint involvement, synovitis, cartilage, and bone erosions. Several studies reported a major role of IL-1β in disease development in CIA mouse model [18], as well as a pathologic role of IL-6 in the effector phase of autoimmune arthritis by promoting bone destruction[19].

CIA mice were daily intraperitoneally injected with CB (at 3 mg/kg (mpK), 10 mpK, and 30 mpK) over a two-week period. As a reference drug, the corticosteroid prednisolone (15 mpK) was also injected in CIA mice. CB treatment did not exert any effect on mouse body weight whatever the dose tested overtime (**Fig. 5A**), indicating no major toxicity of CB in treated mice. While CIA mice displayed elevated levels of circulating IL-1β and IL-6 two weeks after disease onset (**Fig. 5B**), CB treatment resulted in a profound decrease of both cytokines (**Fig. 5B**). Of note, the highest dose of CB (30 mpK) was as efficient as the referenced corticosteroid prednisolone.

**Figure 5.**
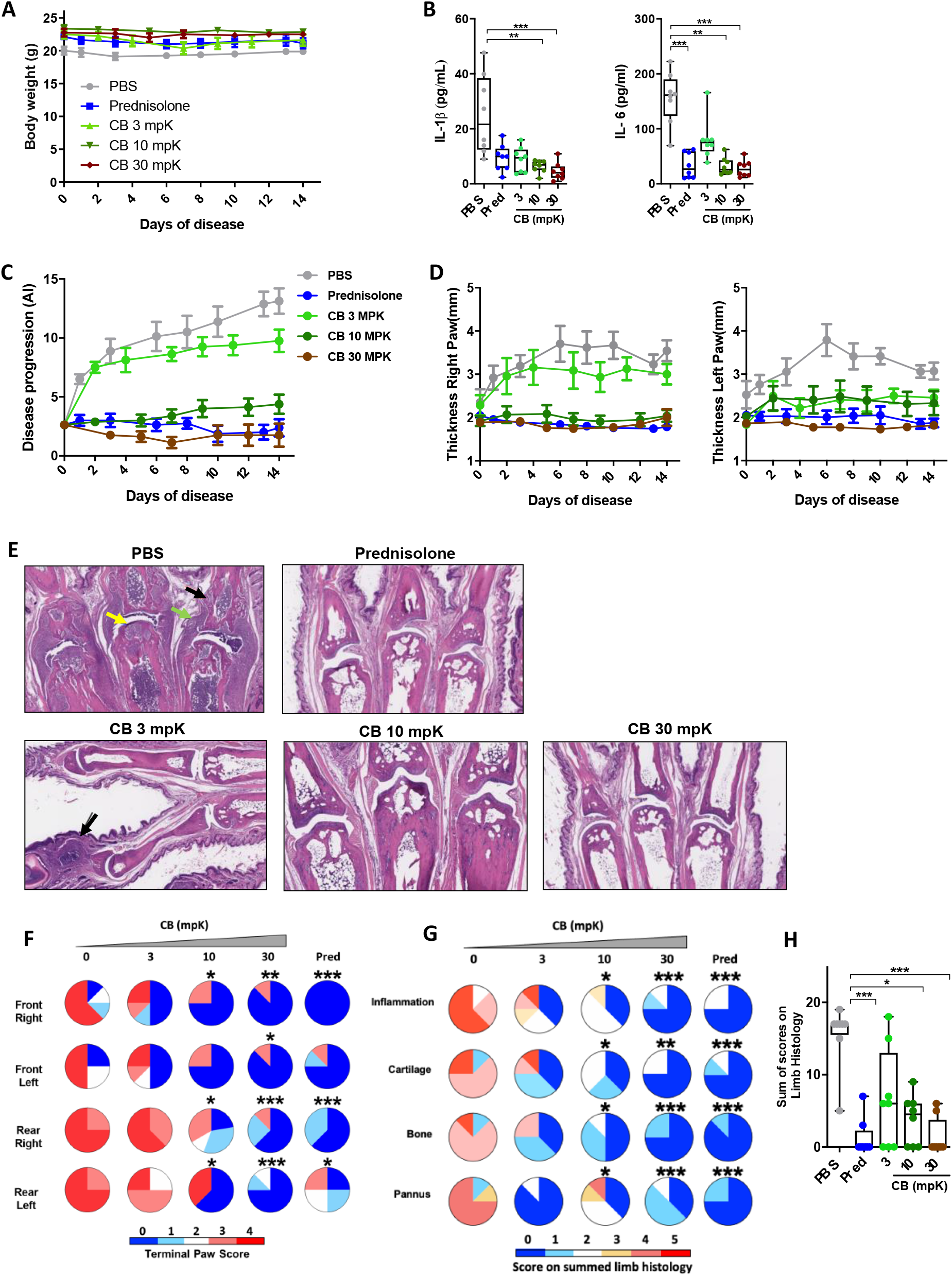
CB controls inflammation and disease onset in collagen induced arthritic mice. (**A**) The body weight of collagen induced arthritic mice treated with prednisolone (Pred, 15 mpK) or 3, 10, or 30 mpK of CB was measured every other day. (**B**) IL-1β and IL-6 were measured by ELISA in the serum after 2 weeks of treatment. Kruskal-Wallis test with Dunn’s post hoc correction. (**C**) the progression of the disease was assessed according to several markers in the legs. (**D**) The thickness of the legs was measured. (**E**) Representative hematoxylin and eosin staining of paw sections from collagen induced arthritic mice treated with prednisolone (Pred, 15 mpK) or 3, 10, or 30 mpK of CB: joints with pannus and inflammation (green arrow), loss of articular cartilage (yellow arrow) and bone remodeling (black arrow). (**F**) Impact of the treatment was evaluated by a paw score, signs of arthritis in all paws according to a 0-4 scale of ascending severity, after 2 weeks of treatments. (**G**) Effects of the treatment on the scoring for inflammation, cartilage damage, bone remodeling, and pannus after 2 weeks of treatment. (**H**) Combined disease score after 2 weeks of treatment. Kruskal-Wallis test with Dunn’s post hoc correction. All data are presented as median ± range, ***P < 0.001, **P < 0.01, *P < 0.05. *n* = 8 mice per group

We also investigated the effect of CB treatment on disease progression. The daily injection of CB attenuated the progression of the disease during the 14 days of treatment (**Fig. 5C**). Mice treated with CB had thinner legs than mice treated with PBS (**Fig.5D**). In arthritic mice, the histological analysis of the tissue injury revealed severe cartilage damage (yellow arrow), bone remodeling (black arrow) and tissue infiltration by immune cells (green arrow) (**Fig. 5E**). Daily CB treatment attenuated all these pathological markers **(Fig. 5E)**. To further characterize the effect of CB treatment, individual paws were scored, reflecting the severity of the disease. CB treatment drastically reduced the terminal paw score (**Fig. 5F)**, which was associated with reduced inflammation, cartilage damage, bone remodeling on limbs and pannus (**Fig. 5G and 5H**).

### CB attenuates spontaneous cytokine production in COVID-19 patients’ whole blood

In COVID-19 severe patients, lung infiltrated monocytes produce high levels of inflammatory cytokines such as IL-1β, IL-6 and TNF-α leading to uncontrolled inflammation[20], “cytokine storm”, and fever[21]. Moreover, COVID-19 patients requiring intensive care in hospitals, exhibit blood plasma levels of proinflammatory cytokines[10], like IL-1β, IL-6, IL-10 and TNF-α [22] higher than HD (**Fig. 6A**). Thus, targeting proinflammatory cytokines and/or monocytes would represent therapeutic strategy for treatment of COVID-19 severe cases. Here, we provide the proof of concept that targeting CXCR4 with CB could represent a novel therapeutic approach for hospitalized COVID-19 patients. According to digital ELISA (Simoa) the treatment of whole blood from COVID-19 patients with CB at 20μM for 16h decreased the spontaneous production of IL-1β, TNF-α, IL-6 and IL-10. (**Fig. 6B**).

**Figure 6.**
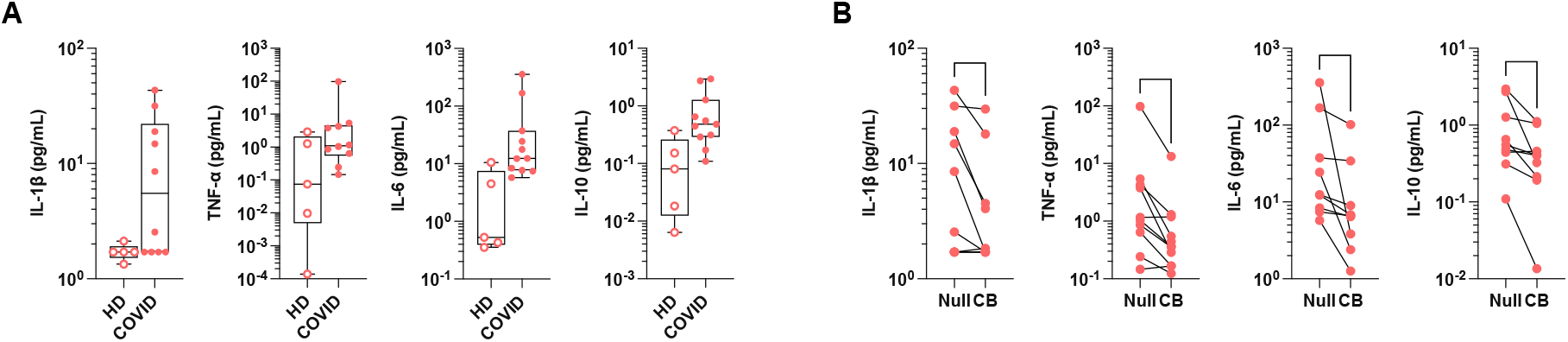
CB controls inflammation in whole blood of COVID-19 patients. (**A**) IL-1β, TNF-α, IL-6 and IL-10 were measured after 16h at 37°C in the plasma of whole blood from healthy donors (HD) and COVID-19 patients by digital ELISA (Simoa). (**B**) Whole blood from HD and COVID-19 patients were treated by CB at 20 μM for 16h and cytokines IL-1β, TNF-α, IL-6 and IL-10 were measured in the supernatant by digital ELISA (Simoa). Box and whisker plots with median ± range. Mann-Whitney test. **P < 0.01, *P < 0.05

## DISCUSSION

In this study, we reveal a broad-spectrum anti-inflammatory activity of the histamine analogue CB. We demonstrate that CB significantly reduces the spontaneous and induced productions of the key inflammatory cytokines by monocytes from blood and synovial fluids of JIA patients. This immuno-modulatory activity of CB strictly depends on the engagement of CXCR4 chemokine receptor. In addition, CB displays high efficiency *in vivo* to reduce inflammation and subsequent tissue damage in arthritic mice, leading to reduction of disease progression. In addition, we provide evidence that targeting CXCR4 with CB also calms hyper inflammation associated with severe COVID-19. Put together, our results suggest that targeting CXCR4 with CB-like molecules represents a promising therapeutic strategy for inflammatory diseases in which hyper activation of monocytes plays a deleterious effect.

The pro-inflammatory cytokines TNF-α and IL-6 are major contributors to JIA and RA [23,24]. Among all pro-inflammatory factors produced during JIA flares, the hierarchical importance of TNF-α and IL-6 is strongly supported by the clinical benefits upon their inhibition. While the TNF-α signature has been clearly identified in JIA patients[25], systemic TNF-α protein remains, however, difficult to detect and measure, notably because most classical ELISAs are not sensitive enough to detect TNF-α concentration below one pg/mL. To overcome this challenge, we developed an ultrasensitive digital TNF-α ELISA (Simoa) with attomolar sensitivity that permits to show that TNF-α concentration in plasma from JIA patients is six-fold higher than the one in healthy donors. Through Simoa experiments, we also measured that TNF-α concentration in synovial fluid JIA patients is higher in their synovial fluid than that in plasma. In most forms of RA, including JIA, blood monocytes are attracted to synovial fluids where they differentiate into inflammatory macrophages[1] and produce joint-degrading mediators. These immune cells emerge as being the major sources of TNF-α. Accordingly, we show that purified monocytes from JIA patients spontaneously produce more TNF-α than monocytes from healthy donors. As previously described by Cepika *et al* [1], we also confirm that monocytes from JIA patients are hyperresponsive to TLR-7/8 activation compared to healthy individuals. In any case, we provide the prime evidence that CB is a potent inhibitor of the production of pro-inflammatory cytokines and chemokines in inflammatory monocytes originating from blood or synovial fluid of JIA patients. Interestingly, CB anti-inflammatory activity is not restricted to TNF-α and IL-6, as the inflammatory signature of JIA patients is largely toned down under CB exposure. These *ex vivo* results suggest that CB potentially reduces hypersecretion of cytokines and chemokines observed in JIA patients during flares. In RA animal models, IL-6 has been reported to stimulate recruitment of leukocytes to inflammatory sites, synoviocyte proliferation and osteoclast activation, resulting in synovial pannus formation[19]. In synergy with IL-1β, IL-6 increases production of matrix metalloproteinases, which contributes to joint and cartilage destruction[26]. In collagen-induced arthritis (CIA) mice, daily treatment of CB exhibits strong anti-inflammatory properties by blocking both IL-1β and IL-6 secretions. According to massive reduction of cartilage damage, bone remodeling, immune cell tissue infiltration and pannus CB treatment results in reduced disease progression and paw thickness in arthritic mice similarly to the referenced corticosteroid prednisolone. These *in vivo* experiments validate the concept of the immunomodulatory activity of CB observed *in vitro* and *ex vivo* on synovial monocytes from JIA patients.

In competition assays using the CXCR4 antagonist AMD3100 or siRNA-based experiments to down-regulate CXCR4 expression, we further demonstrated that CB anti-inflammatory activity on monocytes strictly depends on CXCR4. Beyond the impact of CXCR4 signaling on regulation of the IFN pathway[11], this result highlights that CXCR4 also represents a broad spectrum regulator of inflammation in various cell types, including pDC and monocytes. This CXCR4-mediated dual anti-IFN and anti-inflammatory activity may provide clinical benefits as type I IFNs are widely detected and functionally active in RA[27]. Interestingly, a recent study shows that the levels of CXCR4 and its natural ligand, CXCL12 are significantly higher in the serum and joint synovial fluids of active RA patients compared to the control group[28]. Moreover, CXCR4 and CXCL12 expression levels in the RA active group are higher compared to the remission group[28]. Higher accessibility to CXCR4 in RA patients makes targeting the CXCR4 anti-inflammatory pathway a particularly promising strategy for these pathologies.

For instance, drugs that target TNF-α, IL-6 and IL-1β cytokines or their receptors have shown beneficial effects in JIA patients[23]. However, these treatments are often associated with highly heterogeneous responses across patients in terms of efficacy and treatment resistance. Consequently, it is not uncommon for a patient to change medication throughout the course of the disease. To some extent, this could be explained by the very high specificity of these treatments toward a single cytokine in a set of diseases characterized by a very broad inflammatory spectrum. By contrast to targeted therapies, corticosteroids have been used for decades to block overall pro-inflammation in RA patients, with, however, strong side-effects[29]. To overcome these limitations, novel strategies such as Janus kinases (JAK) inhibitors, which are small molecules inhibiting the activity of JAK, showed reasonable success in adult form of arthritis[30][31][32]. As for strategies targeting JAK, we demonstrate that CB treatment exhibits wide-ranging cytokine inhibition *in vitro, ex vivo* and *in vivo*. CB acts, however, one step upstream of JAK inhibitors, by blocking the production of inflammatory cytokines rather than blocking cytokine-mediated signaling, which may confer some advantages in terms of therapies efficacies. This is in fact supported by the strong inhibition of disease progression in CB-treated RA mice, validating the concept of targeting CXCR4 with CB-like molecules to treat arthritic diseases. CB indeed holds all the required hallmarks for a promising new drug to treat RA: CB is a small molecule, which does not show any side effects in preclinical models *in vivo*, targets the widely expressed receptor CXCR4 on immune cells, and acts on cytokine production with a broad anti-inflammatory spectrum.

To illustrate the potential benefit of CB in another inflammatory disease context, we studied samples from COVID-19 patients. Hypersecretion of inflammatory cytokines by monocytes and macrophages in lung seems to be central in COVID-19 pathology. Studies of SARS-CoV2 infection have revealed highly inflammatory monocyte/macrophage[9,10],[33] population in the bronchoalveolar lavage of patients with severe but not mild COVID-19[34,35]. The lungs infiltrated with monocytes exhibit high levels of inflammatory cytokines such as IL1-β, IL-6[9] and TNF-α at the root of an uncontrolled inflammation[20], a cytokine storm, and fever[21]. We previously showed that patients requiring intensive care in hospitals displayed higher blood plasma levels of IL-6 and TNF-α, which continue to increase over time and correlate with bad prognostic[10]. Therefore, targeting pro-inflammatory cytokines and/or monocytes also represents a potential therapeutic strategy for the treatment of COVID-19 severe cases. Anti-inflammatory treatments have been proposed, based on the use of corticosteroids or JAK inhibitors[36], or the humanized monoclonal anti-IL-6 antibody Tocilizumab. Tocilizumab is FDA-approved for the treatment of several inflammatory disorders, including RA[37]. However, up to now, these commercial drugs have shown conflicting results between studies. We show here, using Simoa, that CB controls the spontaneous productions of IL-1β, TNF-α, IL-6 and IL-10 in whole blood from severe COVID-19 patients. This suggests CB could reduce simultaneously the production of key pro-inflammatory cytokines involved in COVID-19 pathology. Although these results remain preliminary, they nevertheless open the door to the use of drugs targeting CXCR4 as a new therapeutic strategy to control pulmonary hyperinflammation in COVID-19 patients.

In conclusion, the very low cytotoxicity of CB in addition to its broad-spectrum inhibitory activity *ex vivo* on inflammatory cytokine production and its therapeutic activity *in vivo*, suggest that targeting CXCR4 with CB-like molecules could represent a novel promising therapeutic strategy for chronic inflammatory diseases such as rheumatoid arthritis and potentially to acute viral-mediated inflammation such as COVID-19 disease.

## MATERIALS AND METHODS

### Blood samples isolation and culture of blood leukocytes

The blood from healthy donors was obtained from *“Etablissement Français du Sang”* (convention # 07/CABANEL/106), Paris, France. For JIA patients’ material, the study was approved by the Comité de Protection des Personnes (N° EudraCT: 2018-A01358-47) in France. Experimental procedures with human blood were done according to the European Union guidelines and the Declaration of Helsinki and informed consent was obtained from all donors (healthy and patients). The JIA patients’ clinical data are summarized in the Table 1. *In vitro* experiments were performed using human mononuclear cells from peripheral blood or synovial fluid (SF) isolated by centrifugation in density gradient medium (STEMCELL Technologies). Human monocytes were purified by positive selection with Human CD14 microbeads (Miltenyi). Synovial fluid mononuclear cells (SFMC), peripheral blood mononuclear cells (PBMC) and monocytes were cultured in RPMI 1640 (Invitrogen, Gaithersburg, MD) (R10) containing 10% heat-inactivated fetal bovine serum and 1mM glutamine (Hyclone, Logan, UT). The JIA patients were identified from P1 to P36 (Fig. 2).

**Table 1.**
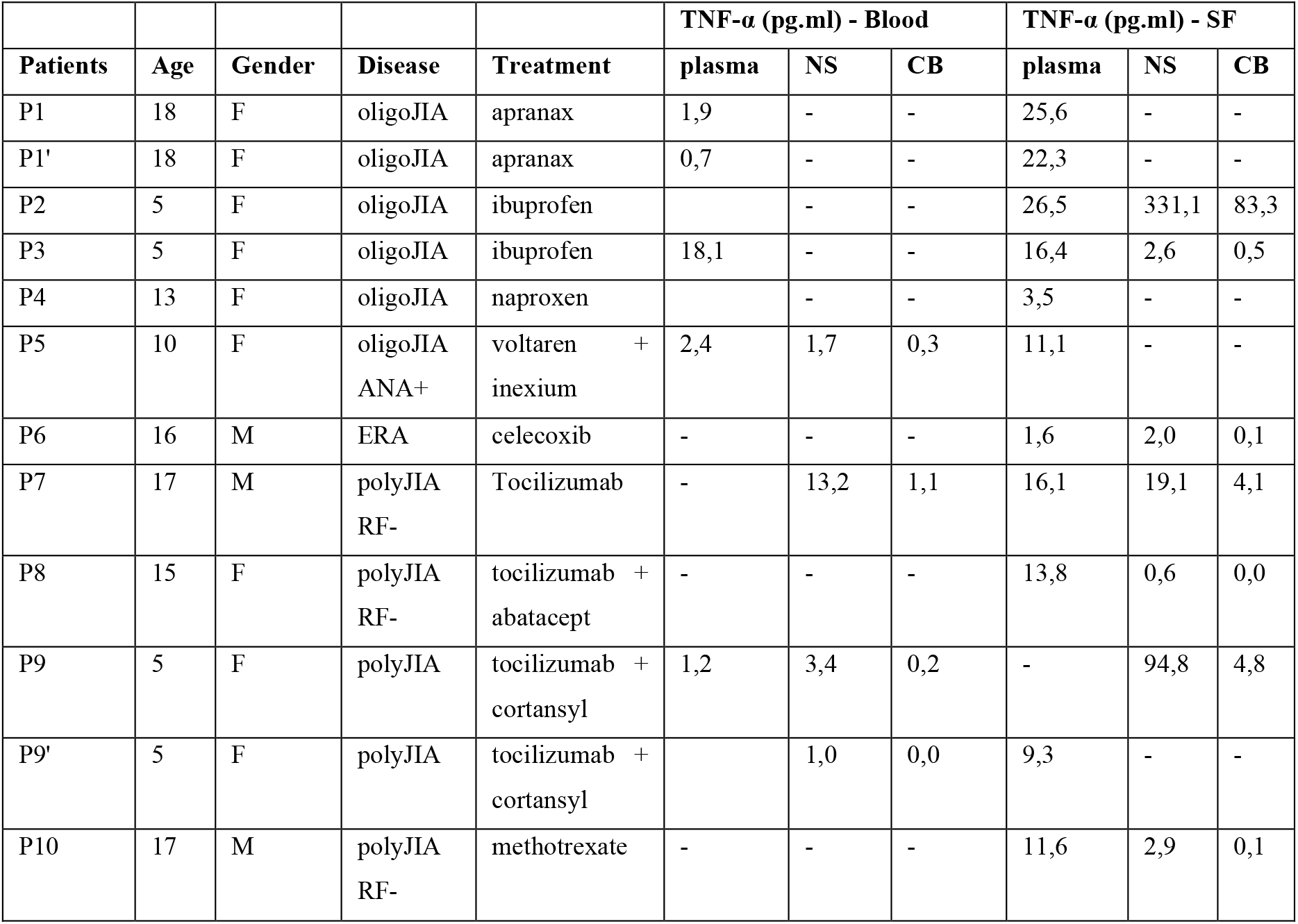
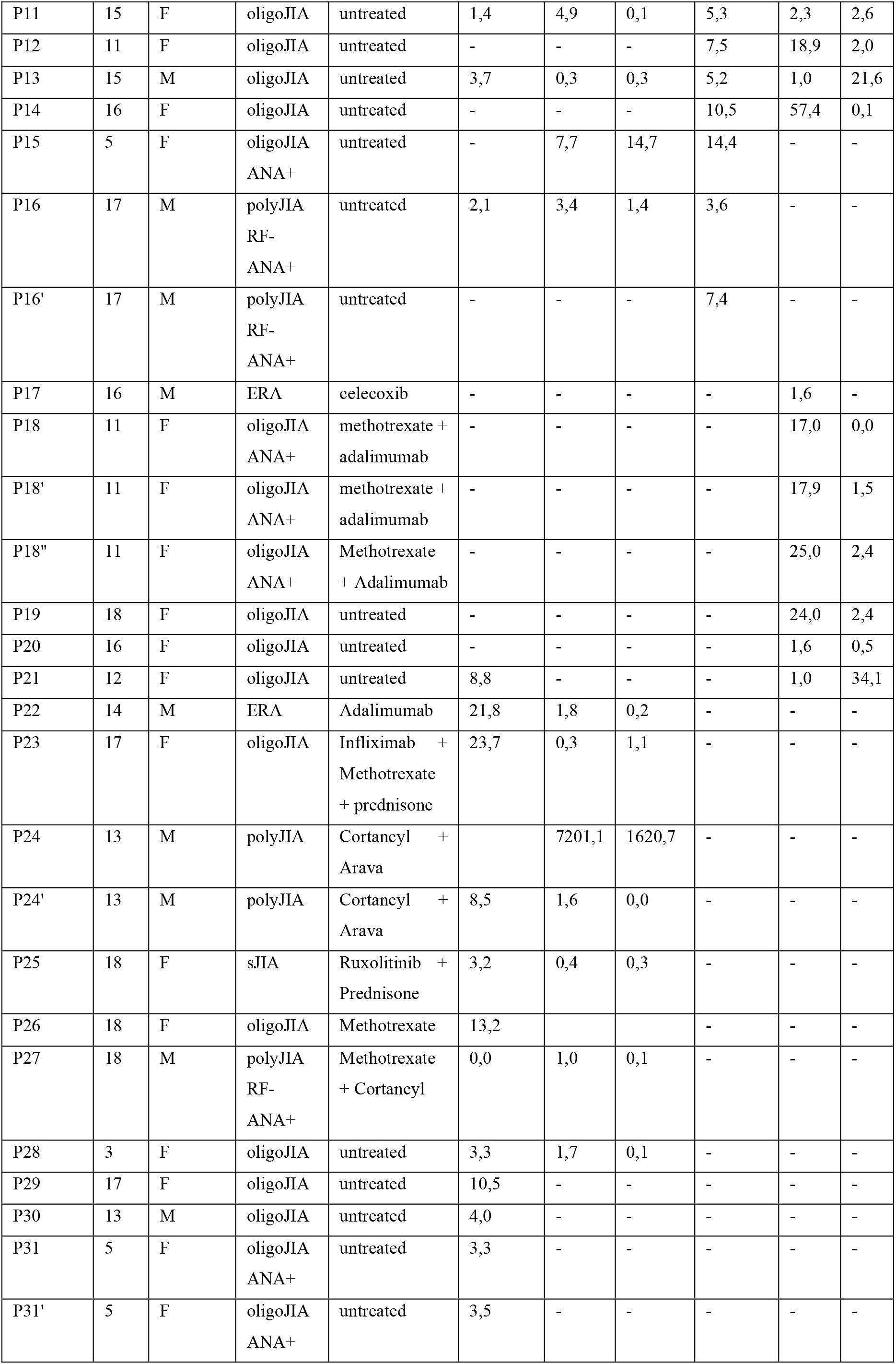
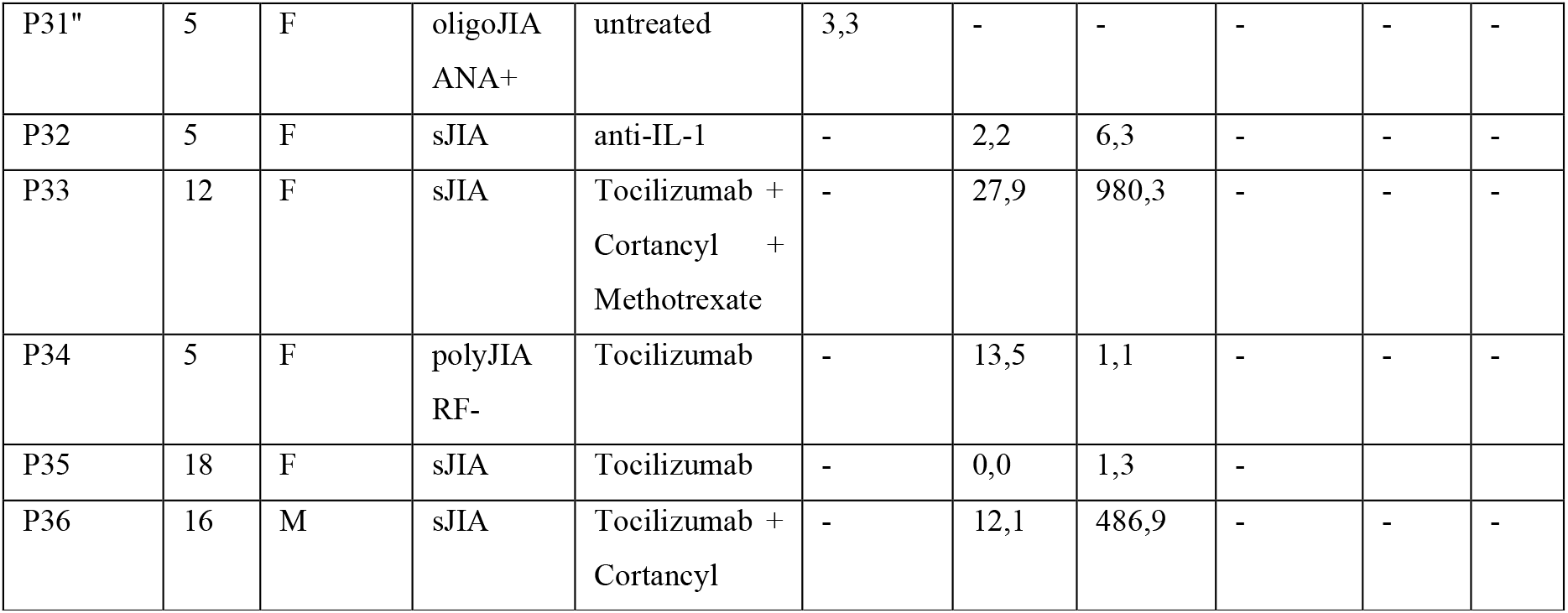
Clinical information of recruited patients. SF: synovial fluid, oligoJIA: oligoarticular juvenile idiopathic arthritis, ERA: enthesitis-related arthritis, polyJIA: polyarticular juvenile idiopathic arthritis, sJIA: systemic juvenile idiopathic arthritis, RF: rheumatoid factor ANA: antinuclear factor

Ten COVID-19 patients were included at the time of their hospitalization in Cochin Hospital. COVID-19 hospitalization is defined by respiratory distress requiring some oxygen ranging from 3L/min of oxygen to mechanical ventilation. Biological collection and informed consent were approved by the Direction de la Recherche Clinique et Innovation (DRCI) and the French Ministry of Research (N°2019-3677). The study conforms to the principles outlined in the Declaration of Helsinki and received approval by the appropriate Institutional Review Board (Cochin-Port Royal Hospital, Paris, France; number AAA-2020-08018).

### Cell stimulation

PBMCs were seeded at 2.10^6^ cells /mL and monocytes isolated from PBMCs or SFMCs were seeded at 1.10^6^ cells /mL. Cells were then stimulated with the TLR-7/8 agonist Resiquimod – R848 (Invivogen) at 5 μg/mL. Cells were pre-treated with AMD3100 (Sigma-Aldrich) and/or clobenpropit (CB) (Sigma Aldrich) at 20 μM (or other if specified) for 1 h prior to 16 h stimulation or as indicated. Whole blood from COVID-19 patients and matched healthy donors were diluted to 1/3 in RPMI 1640 and cultured in the presence or not of CB (20 μM) for 16h. Cells were then collected for flow cytometry or mass cytometry and supernatants were collected for cytokine detection. For intracellular staining, Brefeldin A (BFA) was added to the cells 30 minutes after stimulation for 5 h. RAJOUTER description THP-1 dual.

### Mass Cytometry

PBMCs from healthy donors were stimulated by R848 in the presence of CB and Brefeldin A was added overnight. After stimulation, PBMC were labeled with the cocktail of surface antibodies and Rh isotope in PBS, for 20 min at room temperature (RT). Commercial and in-house-conjugated antibodies were used to create a panel to perform phenotypic and functional analysis. Two DNA interchelator (Rhodium isotope mass 103 and Iridium isotopes mas 191 & 193) were used respectively to determine viability and to gate the singlets cells during analysis. Each antibody has been previously titrated on PBMC. The cells were then fixed for 15 min at RT with Fix-I solution (Maxpar Fix-I buffer; Fluidigm). After fixation, cells were incubated in Perm-S solution containing the anti-cytokine antibodies and isotopes Ir for 30 min at RT (Maxpar Perm-S buffer;Fluidigm). Just before CyTOF acquisition, a series of 3 washes in extremely pure water is carried out and the cells were kept in water at a maximum concentration of 500 000 cells / mL. (Maxpar Water; Fluidigm). Data were acquired using a CyTOF2 instrument (Fluidigm) and analyzed using both FlowJo software for cleaning data and the viSNE algorithm in Cytobank to perform main analysis. The antibodies coupled to the metals are shown in Table 2.

**Table 2.** Antibodies for mass cytometry.

| **Targets** | **Metals** | **Clones** | **Suppliers** | **References** |
| --- | --- | --- | --- | --- |
| CD45 | 89Y | HI30 | Fluidigm | 3089003B |
| CD38 | 112Cd | HIT2 | Life technologies | Q22150 |
| HLA-DR | 115 In | L243 | Ozyme | BLE307651 |
| IL-1b | 141 Pr | 8516 | R & D Systems | MAB201-100 |
| CD27 | 142 Nd | O323 | Ozyme | BLE302802 |
| CD123 | 143 Nd | 6H6 | Ozyme | BLE306002 |
| CD4 | 145 Nd | RPA-T4 | Ozyme | BLE300516 |
| CD8 | 146 Nd | RPT-T8 | Ozyme | BLE301018 |
| CD86 | 148 Nd | FM95 | Miltenyi | 130-095-212 |
| IL-17 | 151 Eu | CZ8-23G1 | Miltenyi | 130-095-212 |
| CD304 (BDCA4) | 152 Sm | AD5-17F6 | Miltenyi | 130-095-212 |
| CD11C | 153 Eu | Bu15 | Ozyme | BLE337202 |
| Il-6 | 154 Sm | MG2-13A5 | Miltenyi | 130-095-212 |
| CD279 (PD-1) | 156 Gd | PD1.3.1.3 | Miltenyi | 130-095-212 |
| CD14 | 160 Gd | M5E2 | Ozyme | BLE301810 |
| IFNγ | 161 Dy | 45-15 | Miltenyi | 130-095-212 |
| CD185 (CXCR5) | 162 Dy | REA103 | Miltenyi | 130-095-212 |
| CCR7 (CD197) | 163 Dy | REA108 | Miltenyi | 130-095-212 |
| IFN α2 | 165 Ho | 3C5 | R & D Systems | H00003440-M27 |
| TNFα | 166 Er | cA2 | Miltenyi | 130-095-212 |
| CD56 | 168 Er | HCD56 | Ozyme | BLE318324 |
| CD45RA | 169 Tm | T6D11 | Miltenyi | 130-095-212 |
| CD3 | 170 Er | BW264/56 | Miltenyi | 130-095-212 |
| IgM | 172 Yb | PJ2-22H3 | Miltenyi | 130-095-212 |
| IgD | 174 Yb | IgD26 | Miltenyi | 130-095-212 |
| CXCL10 | 175 Lu | REA334 | Miltenyi | 130-095-212 |
| CD19 | 176 Yb | LT19 | Miltenyi | 130-108-029 |
| CD278 (ICOS) | 198 Pt | REA183 | Miltenyi | 130-095-212 |
| CD16 | 209 Bi | 3G8 | Fluidigm | 3209002B |

### Flow cytometry

Cells were washed in PBS and then incubated with a viability stain (Zombie-Aqua, Biolegend) for 30 min at 4°C. After washing, the cells were resuspended in PBS containing 2% FCS and 2mM EDTA and stained with the extracellular mix using APC-Vio770 anti-CD14 (clone REA599) from Miltenyi Biotec and used at 1/100. For the TNF-α and IL-1β intracellular staining, an Inside Stain kit (Miltenyi Biotec) was used according to the manufacturer’s protocol. Briefly, the cells were fixed for 20 min at RT with 250 μL of the Inside Fix solution then washed and stained in 100 μL of the Inside Perm solution containing PE anti-TNF (clone cA2, Miltenyi Biotec) and APC anti-IL-1β (clone REA1172, Miltenyi Biotec) antibody at 1/50 for 30 min at room temperature. Data acquisition was performed on a Canto II flow cytometer using Diva software (BD Biosciences, San Jose, CA). FlowJo software (Treestar, Ashland, OR) was used to analyze data.

### CXCR4 knockout experiments

Monocytes were seeded at 10^5^ cells/100 μl in 96-well plates and incubated at 37°C. Control siRNA (qiagen) and CXCR4 siRNA (SMARTPool, Dharmarcon) were diluted in DOTAP (1,2-dioleoyl-3-trimethylammonium-propane; Roche Applied Sciences). The mix was gently mixed and incubated at room temperature for 15 min. After incubation, the mix was added to cells in culture at a final concentration of 160 nM. Last, cells were incubated at 37°C for 24 hours before adding treatments and stimulation.

### Cytokine detection

Supernatants were tested for multiple cytokine production using the bead assay LEGENDplex Antivirus Human panel (Biolegend, San Diego, USA) or with a commercial Luminex multi-analyte assay (Biotechne, R&D systems) according to the manufacturer’s instructions.

### Simoa

Simoa digital ELISA specific for TNF-α was developed using a Quanterix Homebrew Assay and two commercially available antibodies[38]. The 28401 antibody (R&D) clone was used as a capture antibody after coating paramagnetic beads (0.3 mg/mL), and the ab9635 polyclonal antibody (Abcam) was biotinylated (biotin/Ab ratio = 30:1) and used as the detector. Recombinant TNF-α (R&D) was used as a standard curve after cross-reactivity testing. The limit of detection was calculated by the mean value of all blank runs ± 3 SDs and was 1 fg/mL.

### RNA isolation and real-time quantitative RT-PCR analyses

Human THP1-Dual monocytes cells seeded at 1.6.10^6^ cells/mL were pre-treated with clobenpropit (CB) for 1 h prior to 24 h stimulation with R848 at 10 μg/mL. Total RNA was isolated using an E.Z.N.A. kit according to the manufacturer’s instructions (Omega Bio-Tek, USA). The first-strand cDNA synthesis was performed with the Prime Script RT Master Mix kit (Takara Bio Europe, France). Quantitative real-time PCR was performed at 60°C using Takyon ROX SYBR MasterMix (Eurogentec, Belgium) in the CFX384 Touch Real-Time PCR Detection System (Bio-Rad, France). Primers used in RT-qPCR analyses are listed below: *RPL13A*: forward primer, 5′-AACAGCTCATGAGGCTACGG-3′; reverse primer, 5′-TGGGTCTTGAGGACCTCTGT-3′ *IL1β*: forward primer, 5′-CCTGTCCTGCGTGTTGAAAGA-3′; reverse primer, 5′-GGGAACTGGGCAGACTCAAA-3′ *IL6*: forward primer, 5′-GACAGCCACTCACCTCTTCA-3′; reverse primer, 5′-CCTCTTTGCTGCTTTCACAC-3′ *TNFα*: forward primer, 5′-CCTGCTGCACTTTGGAGTGA-3′; reverse primer, 5′-GAGGGTTTGCTACAACATGGG-3′

### Nanostring gene expression analysis

Total RNA was extracted from isolated monocytes of controls and patients and was diluted with ribonuclease-free water at 20 ng/μl, and 100 ng (5 μl) of each sample was analyzed using the Human Immunology kit v2 and Nanostring Counter. Each sample was analyzed in a separate multiplexed reaction including, in each, eight negative probes and six serial concentrations of positive control probes. Data were imported into nSolver analysis software (version 2.5) for quality checking and normalization of data according to NanoString analysis guidelines, using positive probes and housekeeping genes. For analysis, mRNA expression levels were log-transformed before hierarchical clustering (Qlucore Omics Explorer version 3.1). Pathway analysis was performed using DAVID bioinformatics databank (https://david.ncifcrf.gov).

### Mice

Animal experiments were performed blindly by Washington Biotechnology, INC. The animals were housed in a temperature and humidity-controlled room with 12-hour light/dark cycles with food and water *ad libitum*. The WBI Animal Care and Use Committee reviewed and approved all mouse experiments (IACUC NO. 17-006). Animal welfare and experimental protocols were strictly complied with the Guide for the Care and Use of Laboratory Animals and the ethics and regulations.

At Day 0, the mice were weighed, hind limb thickness recorded by digital caliper and injected subcutaneously at the base of the tail with 50 μL of the collagen/ Complete Freund’s Adjuvant emulsion. At Day 21, the mice were weighed and injected subcutaneously at the base of the tail with 50 μL of the collagen/ Incomplete Freund’s Adjuvant emulsion, then divided in 5 groups of 8 mices. From Day 21 to day 35, group 1 was treated with intraperitoneal injection, of PBS. All mice in group 2 were treated daily by PO (oral gavage) with 10 mL/kg (mpk), of prednisolone (Sigma, P4153), that was previously dissolved in 3 mL of PBS to prepare a 1.5 mg/mL solution. Mice from group 3, 4 and 5 were treated with CB. Briefly, CB was dissolved in PBS and intraperitoneally injected daily at 3 mg/kg (mpK), 10 mpK or 30 mpK (group 3 to 5). At Day 35, all mice were weighed, scored for signs of arthritis, hind paw volumes recorded. The mice were anesthetized and exsanguinated into pre-chilled EDTA tubes. The blood was processed to plasma, frozen and then was thawed at RT, diluted 1:2, and assayed by ELISA for IL-1β (R&D Systems, Cat. MLB00C) and IL-6 (R&D Systems, Cat. M6000B). The limbs were individually removed to 10% neutral buffered formalin.

### Histology on mice limbs

Formalin-fixed mouse paws were processed routinely, sectioned at approximately 8 microns, and stained with hematoxylin and eosin. Glass slides were evaluated using light microscopy by a board-certified veterinary pathologist. The severity of histologic findings was scored using the following scoring criteria as adapted from Crissman *et al*[39].

- Inflammation 0=Normal 1=Minimal infiltration of inflammatory cells in synovium and periarticular tissue of affected joints 2=Mild infiltration, if paws, restricted to affected joints 3=Moderate infiltration with moderate edema, if paws, restricted to affected joints 4=Marked infiltration affecting most areas with marked edema 5=Severe diffuse infiltration with severe edema
- Cartilage Damage 0=Normal 1=Minimal=minimal to mild loss of toluidine blue staining with no obvious chondrocyte loss or collagen disruption in affected joints 2=Mild=mild loss of toluidine blue staining with focal mild (superficial) chondrocyte loss and/or collagen disruption in affected joints 3=Moderate=moderate loss of toluidine blue staining with multifocal moderate (depth to middle zone) chondrocyte loss and/or collagen disruption in affected joints 4=Marked=marked loss of toluidine blue staining with multifocal marked (depth to deep zone) chondrocyte loss and/or collagen disruption in most joints 5=Severe =severe diffuse loss of toluidine blue staining with multifocal severe (depth to tide mark) chondrocyte loss and/or collagen disruption in all joints
- Bone Resorption 0=Normal 1=Minimal=small areas of resorption, not readily apparent on low magnification, rare osteoclasts in affected joints 2=Mild=more numerous areas of, not readily apparent on low magnification, osteoclasts more numerous in affected joints 3=Moderate=obvious resorption of medullary trabecular and cortical bone without full thickness defects in cortex, loss of some medullary trabeculae, lesion apparent on low magnification, osteoclasts more numerous in affected joints 4=Marked=Full thickness defects in cortical bone, often with distortion of profile of remaining cortical surface, marked loss of medullary bone, numerous osteoclasts, affects most joints 5=Severe=Full thickness defects in cortical bone and destruction of joint architecture of all joints

## STATISTICS

Cellular data sets were analyzed by Mann-Whitney test to compare the group with CB treatment to the untreated group. Flow cytometry data were performed using FlowJo software.

Animal data are shown as median and min/max. Data sets were analyzed by ANOVA (Kruskal-Wallis) tests with Dunn’s post-test for multiple comparisons with the different concentrations of CB and prednisolone treatment. GraphPad Prism 8 (GraphPad Software, San Diego, CA) was used for data analysis and preparation of all graphs. P-values less than 0.05 were statistically significant. Heatmaps were generated with Qlucore OMICS explore Version 3.5(26).

## ACKNOWEDGMENTS

We thank Stephanie Dupuy from the Common Service of Flux Cytometry (B2M) of Paris Descartes University. We also thank Fabrice Porcheray for his help in mass cytometry at BIOASTER Paris.

## FOUNDERS

This work was supported by the Agence National de la Recherche sur le SIDA et les Hépatites ANRS (to J.-P.H.) for the experiments and N.B. Fellowship (AAP 2017-166). NS is a recipient of the European Molecular Biology Organization (EMBO) Long-Term Fellowship (LT-834-2017) and the Pasteur-Roux-Cantarini Fellowship.

## AUTHOR CONTRIBUTORS

N.B, N.S, M.P.R and J.-P.H conceived and designed the study and had full access to all of the data in the study and take responsibility for the integrity of the data and the accuracy of the data analysis. N.B, N.S., P.Q, and M.P.R contributed to writing the paper; and B.S. and J.-P.H wrote the paper. N.B, N.S, B.B, T.D, A.B and J.-P.H performed data analysis. R.M, S.B, B.B.M, B.T and P.Q, were involved in the clinical study and sample collection. B.B, A.L, T.D, A.B, S.N, and M.P.R performed specific experiments and/or analysis. All authors reviewed the manuscript and gave final approval for the version to be published. All authors agree to be accountable for all aspects of the work in ensuring that questions related to the accuracy or integrity of any part of the work are appropriately investigated and resolved.

## Data Availability

The datasets generated during and/or analyzed during the current study are available from the corresponding author on reasonable request.

## REFERENCES

1. Szekanecz Z, Koch AE. Macrophages and their products in rheumatoid arthritis. Current Opinion in Rheumatology 2007;19:289–95. doi:10.1097/BOR.0b013e32805e87ae

2. Yeo L, Adlard N, Biehl M, et al. Expression of chemokines CXCL4 and CXCL7 by synovial macrophages defines an early stage of rheumatoid arthritis. Ann Rheum Dis 2016;75:763–71. doi:10.1136/annrheumdis-2014-206921

3. Petty RE, Southwood TR, Manners P, et al. International League of Associations for Rheumatology classification of juvenile idiopathic arthritis: second revision, Edmonton, 2001. J Rheumatol 2004;31:390–2.

4. Martini A, Lovell DJ. Juvenile idiopathic arthritis: state of the art and future perspectives. Ann Rheum Dis 2010;69:1260–3. doi:10.1136/ard.2010.133033

5. Ou L-S, See L-C, Wu C-J, et al. Association between Serum Inflammatory Cytokines and Disease Activity in Juvenile Idiopathic Arthritis. Clin Rheumatol 2002;21:52–6. doi:10.1007/s100670200012

6. Möller B, Villiger PM. Inhibition of IL-1, IL-6, and TNF-α in immune-mediated inflammatory diseases. Springer Semin Immun 2006;27:391. doi:10.1007/s00281-006-0012-9

7. Mangge H, Kenzian H, Gallistl S, et al. Serum cytokines in juvenile rheumatoid arthritis. Correlation with conventional inflammation parameters and clinical subtypes. Arthritis Rheum 1995;38:211–20. doi:10.1002/art.1780380209

8. Cepika A-M, Banchereau R, Segura E, et al. A multidimensional blood stimulation assay reveals immune alterations underlying systemic juvenile idiopathic arthritis. J Exp Med 2017;214:3449–66. doi:10.1084/jem.20170412

9. Conti P, Ronconi G, Caraffa A, et al. Induction of pro-inflammatory cytokines (IL-1 and IL-6) and lung inflammation by Coronavirus-19 (COVI-19 or SARS-CoV-2): anti-inflammatory strategies. J Biol Regul Homeost Agents 2020;34. doi:10.23812/CONTI-E

10. Hadjadj J, Yatim N, Barnabei L, et al. Impaired type I interferon activity and inflammatory responses in severe COVID-19 patients. Science Published Online First: 13 July 2020. doi:10.1126/science.abc6027

11. Smith N, Pietrancosta N, Davidson S, et al. Natural amines inhibit activation of human plasmacytoid dendritic cells through CXCR4 engagement. Nat Commun 2017;8:14253. doi:10.1038/ncomms14253

12. Gualtierotti R, Ingegnoli F, Griffini S, et al. Prothrombotic biomarkers in patients with rheumatoid arthritis: the beneficial effect of IL-6 receptor blockade. Clin Exp Rheumatol 2016;34:451–8.

13. Huang DW, Sherman BT, Lempicki RA. Systematic and integrative analysis of large gene lists using DAVID bioinformatics resources. Nat Protoc 2009;4:44–57. doi:10.1038/nprot.2008.211

14. Huang DW, Sherman BT, Lempicki RA. Bioinformatics enrichment tools: paths toward the comprehensive functional analysis of large gene lists. Nucleic Acids Res 2009;37:1–13. doi:10.1093/nar/gkn923

15. de Jager W, Hoppenreijs EPAH, Wulffraat NM, et al. Blood and synovial fluid cytokine signatures in patients with juvenile idiopathic arthritis: a cross-sectional study. Ann Rheum Dis 2007;66:589–98. doi:10.1136/ard.2006.061853

16. Km P, M J, B P, et al. Collagen-Induced Arthritis: A model for Murine Autoimmune Arthritis. Bio Protoc 2015;5. doi:10.21769/bioprotoc.1626

17. Hegen M, Keith JC, Collins M, et al. Utility of animal models for identification of potential therapeutics for rheumatoid arthritis. Ann Rheum Dis 2008;67:1505–15. doi:10.1136/ard.2007.076430

18. Williams RO, Marinova-Mutafchieva L, Feldmann M, et al. Evaluation of TNF-α and IL-1 Blockade in Collagen-Induced Arthritis and Comparison with Combined Anti-TNF-α/Anti-CD4 Therapy. The Journal of Immunology 2000;165:7240–5. doi:10.4049/jimmunol.165.12.7240

19. Lipsky PE. Interleukin-6 and rheumatic diseases. Arthritis Res Ther 2006;8 Suppl 2:S4. doi:10.1186/ar1918

20. Blanco-Melo D, Nilsson-Payant BE, Liu W-C, et al. Imbalanced Host Response to SARS-CoV-2 Drives Development of COVID-19. Cell 2020;181:1036–1045.e9. doi:10.1016/j.cell.2020.04.026

21. Merad M, Martin JC. Pathological inflammation in patients with COVID-19: a key role for monocytes and macrophages. Nature Reviews Immunology 2020;20:355–62. doi:10.1038/s41577-020-0331-4

22. Huang C, Wang Y, Li X, et al. Clinical features of patients infected with 2019 novel coronavirus in Wuhan, China. The Lancet 2020;395:497–506. doi:10.1016/S0140-6736(20)30183-5

23. McInnes IB, Buckley CD, Isaacs JD. Cytokines in rheumatoid arthritis - shaping the immunological landscape. Nat Rev Rheumatol 2016;12:63–8. doi:10.1038/nrrheum.2015.171

24. Feldmann M, Brennan FM, Maini RN. Rheumatoid Arthritis. Cell 1996;85:307–10. doi:10.1016/S0092-8674(00)81109-5

25. Prince FH, Dorai Raj AK, Otten MH, et al. TNF-alpha inhibitors for juvenile idiopathic arthritis. Cochrane Database Syst Rev 2018;2018:CD008598. doi:10.1002/14651858.CD008598.pub2

26. Wong PKK, Campbell IK, Egan PJ, et al. The role of the interleukin-6 family of cytokines in inflammatory arthritis and bone turnover. Arthritis Rheum 2003;48:1177–89. doi:10.1002/art.10943

27. Rönnblom L, Eloranta M-L. The interferon signature in autoimmune diseases. Curr Opin Rheumatol 2013;25:248–53. doi:10.1097/BOR.0b013e32835c7e32

28. Peng L, Zhu N, Mao J, et al. Expression levels of CXCR4 and CXCL12 in patients with rheumatoid arthritis and its correlation with disease activity. Experimental and Therapeutic Medicine 2020;20:1925–34. doi:10.3892/etm.2020.8950

29. Hua C, Buttgereit F, Combe B. Glucocorticoids in rheumatoid arthritis: current status and future studies. RMD Open 2020;6:e000536. doi:10.1136/rmdopen-2017-000536

30. Fleischmann R, Mysler E, Hall S, et al. Efficacy and safety of tofacitinib monotherapy, tofacitinib with methotrexate, and adalimumab with methotrexate in patients with rheumatoid arthritis (ORAL Strategy): a phase 3b/4, double-blind, head-to-head, randomised controlled trial. Lancet 2017;390:457–68. doi:10.1016/S0140-6736(17)31618-5

31. Fleischmann R, Kremer J, Cush J, et al. Placebo-controlled trial of tofacitinib monotherapy in rheumatoid arthritis. N Engl J Med 2012;367:495–507. doi:10.1056/NEJMoa1109071

32. Vieira M-C, Zwillich SH, Jansen JP, et al. Tofacitinib Versus Biologic Treatments in Patients With Active Rheumatoid Arthritis Who Have Had an Inadequate Response to Tumor Necrosis Factor Inhibitors: Results From a Network Meta-analysis. Clin Ther 2016;38:2628–2641.e5. doi:10.1016/j.clinthera.2016.11.004

33. Tay MZ, Poh CM, Rénia L, et al. The trinity of COVID-19: immunity, inflammation and intervention. Nature Reviews Immunology 2020;:1–12. doi:10.1038/s41577-020-0311-8

34. Liao M, Liu Y, Yuan J, et al. The landscape of lung bronchoalveolar immune cells in COVID-19 revealed by single-cell RNA sequencing. medRxiv 2020;:2020.02.23.20026690. doi:10.1101/2020.02.23.20026690

35. Zhou Y, Fu B, Zheng X, et al. Pathogenic T cells and inflammatory monocytes incite inflammatory storm in severe COVID-19 patients. Natl Sci Rev Published Online First: 13 March 2020. doi:10.1093/nsr/nwaa041

36. Zhang W, Zhao Y, Zhang F, et al. The use of anti-inflammatory drugs in the treatment of people with severe coronavirus disease 2019 (COVID-19): The Perspectives of clinical immunologists from China. Clin Immunol 2020;214:108393. doi:10.1016/j.clim.2020.108393

37. Nakahara H, Nishimoto N. Anti-interleukin-6 receptor antibody therapy in rheumatic diseases. Endocr Metab Immune Disord Drug Targets 2006;6:373–81. doi:10.2174/187153006779025694

38. Wu D, Milutinovic MD, Walt DR. Single molecule array (Simoa) assay with optimal antibody pairs for cytokine detection in human serum samples. Analyst 2015;140:6277–82. doi:10.1039/c5an01238d

39. Crissman JW, Goodman DG, Hildebrandt PK, et al. Best practices guideline: toxicologic histopathology. Toxicol Pathol 2004;32:126–31. doi:10.1080/01926230490268756

